# Rigid monoclonal antibodies improve detection of SARS-CoV-2 nucleocapsid protein

**DOI:** 10.1101/2021.01.13.426597

**Authors:** Curtis D. Hodge, Daniel J. Rosenberg, Mateusz Wilamowski, Andrzej Joachimiak, Greg L. Hura, Michal Hammel

## Abstract

Monoclonal antibodies (mAbs) are the basis of treatments and diagnostics for pathogens and other biological phenomena. We conducted a structural characterization of mAbs against the N-terminal domain of nucleocapsid protein (NP^NTD^) from SARS-CoV-2 using small angle X-ray scattering (SAXS). Our solution-based results distinguished the mAbs’ flexibility and how this flexibility impacts the assembly of multiple mAbs on an antigen. By pairing two mAbs that bind different epitopes on the NP^NTD^, we show that flexible mAbs form a closed sandwich-like complex. With rigid mAbs, a juxtaposition of the Fabs is prevented, enforcing a linear arrangement of the mAb pair, which facilitates further mAb polymerization. In a modified sandwich ELISA, we show the rigid mAb-pairings with linear polymerization led to increased NP^NTD^ detection sensitivity. These enhancements can expedite the development of more sensitive and selective antigen-detecting point-of-care lateral flow devices (LFA), key for early diagnosis and epidemiological studies of SARS-CoV-2 and other pathogens.

## Introduction

SARS-CoV-2 nucleocapsid proteins (NP) are key for incorporating and packaging viral genomic RNA into mature virions. In infected cells, NPs are produced in large amounts from subgenomic mRNA and are present at the replication-transcription complexes (RTCs), the sites of RNA synthesis. The NP gene is relatively conserved with a sequence identity of 91% and 50% to SARS-CoV and MERS-CoV, respectively, and is rather stable as it acquires few mutations over time (Marra et al., 2003; Zhu et al., 2005). Although the NP from SARS-CoV-2 is abundant and highly immunogenic (Hachim et al., 2020; Ng et al., 2020; Wolfel et al., 2020), most SARS-CoV-2 detection assays use different regions of the spike protein as the antigen in immunoassays. This is mainly because antibodies against the spike protein are believed to be less cross-reactive (Huang et al., 2020) and are expected to correlate better with neutralizing capacity (Poh et al., 2020). Testing for serum antibodies against NP from SARS-CoV-2 was suggested as a way to increase diagnostic capacity (Liu et al., 2020; Ng et al., 2020; Rikhtegaran Tehrani et al., 2020). However, serological assays cannot achieve diagnosis early in the onset of infection as seroconversion occurs after 7-10 days in patients (Jiang et al., 2020; Ng et al., 2020; Wolfel et al., 2020).

Direct detection of viral proteins, often referred to as antigen-based detection, is more sensitive than serology assays in the case of SARS-CoV (Di et al., 2005). An added advantage of antigen-based detection is they are amenable to rapid point-of-care lateral flow assays (LFA). Thus far, antigen-based LFAs are significantly less sensitive than gold-standard RT-PCR but, may approach RT-PCR’s clinical sensitivity with further research and development. The choice of antigen, mAbs, and LFA protocols remain to be fully optimized for SARS-CoV-2.

The abundance and structure of NP in each virion provides a detection advantage over other antigen targets. NP is a 422 amino acid, 46 kDa phosphoprotein composed of two domains linked via a Ser/Arg rich linker with a short C-terminal region. NP dimerizes through its C-terminal domain (CTD) (Zeng et al., 2020). The N-terminal domain (NP^NTD^) is exposed and interacts with RNA. The independent NP^NTD^ and CTD domains do not have stable tertiary contacts in the absence of RNA (Chang et al., 2006; Zeng et al., 2020). In the presence of RNA, NP^NTD^ and CTD form a single bipartite RNA interaction site, which constitutes the basic building block of the nucleocapsid of SARS-CoV-2 (Peng et al., 2020; Yao et al., 2020). Abundance, stability (Zeng et al., 2020), and the location at the surface of higher-order ribonucleoprotein assembly on the RNA (Gui et al., 2017; Yao et al., 2020), makes the NP^NTD^ a viable antigen for the selection of highly specific mAbs for functional assays. NP is one of the early diagnostic markers in SARS-CoV-2 (Li et al., 2020) and has been detected one day before the onset of clinical symptoms in SARS infections (Che et al., 2004). Diagnostic fluorescence LFA immunoassays have been developed to detect SARS-Cov-2 NP protein in nasopharyngeal and nasal swab specimens (Diao et al., 2020; Grant et al., 2020).

LFA protocols could take advantage of antibody-mediated antigen-dependent aggregation into large particles, termed agglutination (Murphy, 2008). These are influenced by antigen valency, enhancing antigen-antibody complex formation (Knutson, van Es, Kayser, & Glassock, 1979; van Es, Knutson, Kayser, & Glassock, 1979). Agglutination is also a factor when pairs of mAbs are used. LFAs that rely on a pair of mAbs that interact with different epitopes on an antigen have improved LFA sensitivity and specificity (Qiu et al., 2005). MAb-NP agglutination can serve to enhance the antigen-based detection limits against NP.

IgG flexibility, its importance in improving mAb recognition, and its influence on agglutination have remained uncharacterized. Although there have been several attempts by cryo-electron tomography (Bongini et al., 2005; Jay et al., 2018; Lei et al., 2019; Sandin, Ofverstedt, Wikstrom, Wrange, & Skoglund, 2004), and negative stain electron tomography (X. Zhang et al., 2015), large scale flexibility measurements are often not amenable to single-particle techniques. In contrast, the resolution of SAXS is sufficient (especially when atomic structures of individual components are available) to determine the Fab regions’ conformational variability in various antibodies, including complexes with antigens or Fc-gamma receptors (FcγRs) (Wright, Elliston, Hui, & Perkins, 2019; Yanaka, Yogo, & Kato, 2020). A previous study showed that Fab regions’ conformational flexibility is derived from the Fc regions’ inherent plasticity in solution (Remesh, Armstrong, Mahan, Luo, & Hammel, 2018).

Here, we used SAXS and other biophysical techniques to structurally characterize mAbs that specifically bind the minimal NP^NTD^ region from a pool of 9 commercial mAbs raised against full-length NP. We correlated the observed flexibilities with super-structures formed when mAb pairs bind NP^NTD^. Our structural insights have general implications for all antigen-antibody interactions. Simultaneously, a novel ELISA assay protocol is intended to expedite the development of sensitive and selective antigen detecting LFAs, which could be applied in early diagnosis and epidemiological studies of SARS-CoV-2.

## Results

### mAbs against nucleocapsid N-terminal domain (NP^NTD^)

We employed an integrative approach by size exclusion chromatography (SEC) coupled with SAXS and multi-angle light scattering (SEC-MALS-SAXS) to find mAbs that selectively bind minimal NP^NTD^. SEC-MALS-SAXS experiments show that from the pool of nine commercial mAbs raised against full-length NP, four antibodies (mAb1, mAb2, mAb4, and mAb8) bind NP^NTD^. The SEC signal shifts with an increase in molecular weight (Figure 1A, Table 1), which shows that mAb1, mAb2, mAb4, and mAb8 form complexes with the NP^NTD^ in a 1:2 molar ratio. Additionally, the radius of gyration (Rg) values distinguish binder from non-binders (Figure 1B, Table 1). Final merged SAXS profiles for the corresponding SEC peak (Supplemental Figure 1) were used to calculate pair-distribution functions (P(r)).

**Figure 1.**
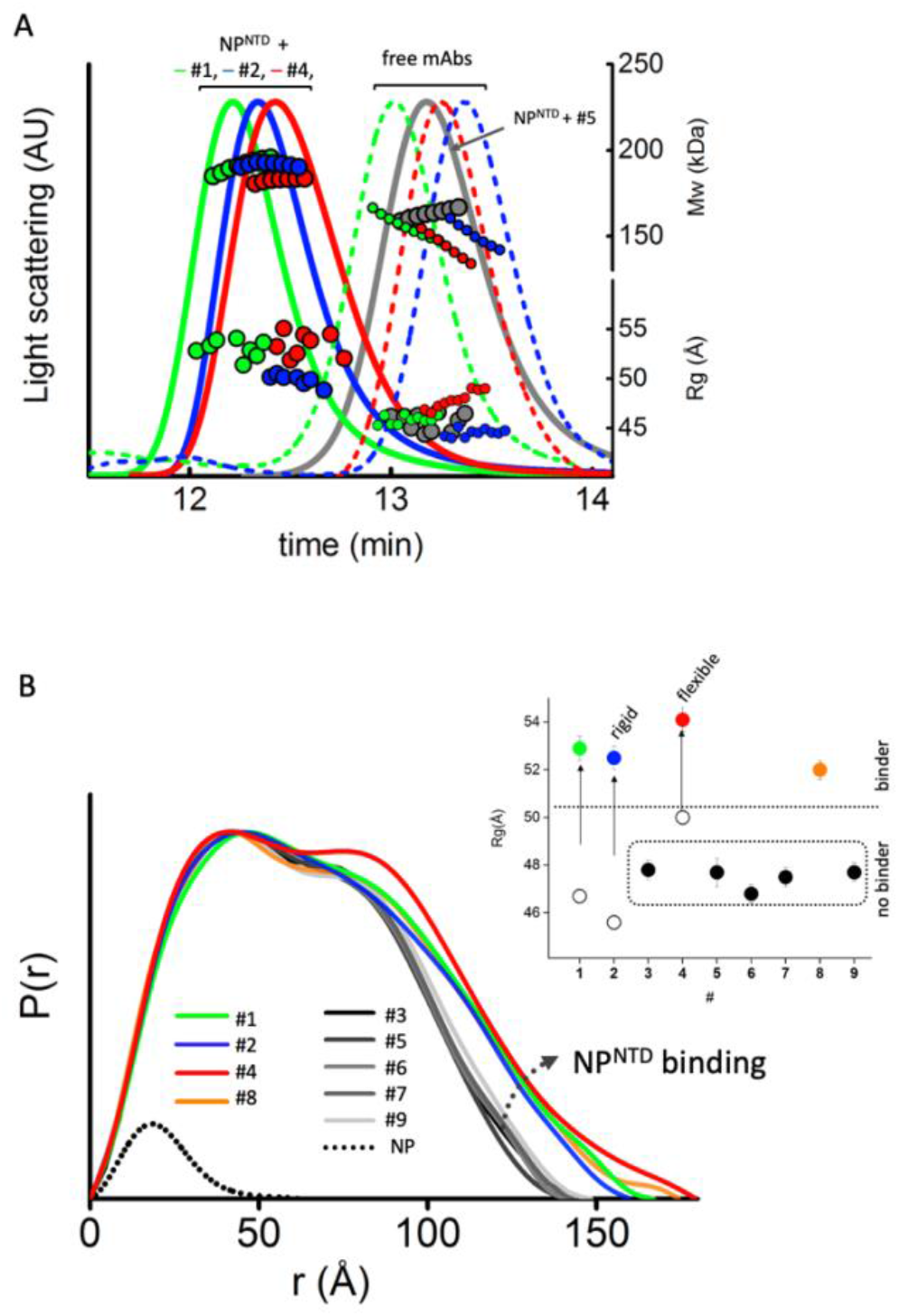
SEC-MALS-SAXS identifies mAbs that bind to NP^NTD^. **A)** SEC-MALS-SAXS chromatograms for free and NP^NTD^ bind mAb1, 2, and 4 (green, blue and red lines). Chromatogram for mAb3 + NP^NTD^ (gray) sample is included for the comparison of the no-binder. Solid lines represent the light scattering signal in arbitrary units, while symbols represent molecular mass (top) calculated from MALS and Rg values (bottom) for each collected SAXS frame versus elution time. **B)** P(r) functions calculated for the experimental SAXS curves for all tested mAb + NP^NTD^ samples (colored as indicated). The P(r) functions are normalized at the maxima. The experimental P(r) function for NP^NTD^ alone is shown for the comparison and normalized relative to the MW estimated by SAXS (Rambo & Tainer, 2013). Inset: Experimental Rg values determined by Guinier plot for the experimental SAXS curves of mAb+NP^NTD^ mixtures (solid dots) and mAb1, 2, and 4 (circles) indicate binder and no-binder. Experimental SAXS curves for mAbs + NP^NTD^ and free mAb1, 2, and 4 are shown in Supplemental Figure 1 and Figure 2B, respectively.

**Table 1.** SAXS, MALS, and SPR experimental parameters

|  | mAbs<br>(#) | R<sub>g</sub><br>(Å) | D<sub>max</sub> | MW<br>MALS/SAXS<br>(kDa) | K<sub>D</sub><br>mAbs/+HRP<br>(pM) |
| --- | --- | --- | --- | --- | --- |
| + NP<sup>NTD</sup> | 1 | 52.9±0.5 | 165 | 193/177 | 1.3/11 |
|  | 2 | 52.5±0.5 | 160 | 190/173 | 190 |
|  | 3 | 47.8±0.4 | 145 | 152/145 |  |
|  | 4 | 54.1±0.5 | 180 | 184/170 | 11/28 |
|  | 5 | 47.7±0.6 | 145 | 162/141 |  |
|  | 6 | 46.8±0.4 | 145 | 154/144 |  |
|  | 7 | 47.5±0.4 | 145 | 152/143 |  |
|  | 8 | 52.0±0.4 | 175 | 184/160 |  |
|  | 9 | 47.7±0.4 | 150 | 170/144 |  |
| free mAbs | 1 | 46.7±0.3 | 145 | 158/157 |  |
|  | 2 | 45.6±0.3 | 140 | 150/147 |  |
|  | 4 | 50.0±0.3 | 155 | 150/167 |  |
| No-<br>pair | 1-4<br>+NP<sup>NTD</sup> | 53.2±0.5 | 180 | 190/169 |  |
| Pair | 1-2<br>+NP<sup>NTD</sup> | 73.9±0.9 | ~300 | 380/397 |  |
|  | 2-4<br>+NP<sup>NTD</sup> | 67.6±0.6 | ~280 | 340/320 |  |

MAb binding of antigen is clearly distinguished by broad P(r) functions relative to those that remain unbound. The P(r) shape further provides information on the overall arrangement of mAb-antigen complexes (Figure 1B), which can be linked to the Fab region’s flexibility (Figure 2A). The first peaks in the P(r) function at r ∼40Å arise from the approximate repeated distances across the Fc or Fab regions’ length and breadth. The P(r) shoulder at r∼80Å reflects the inter-domain distances between Fc and Fab regions. Simultaneously, the divide between P(r) peak and shoulder reflects Fab regions’ distancing that correlates with the extended conformers’ occupancy in solution (Wright et al., 2019). The P(r) features and experimental Rg values (Figure 1B) allowed us to rank the inherent flexibility of mAbs, with mAb2 adopting the least and mAb4 the most extended states.

**Figure 2.**
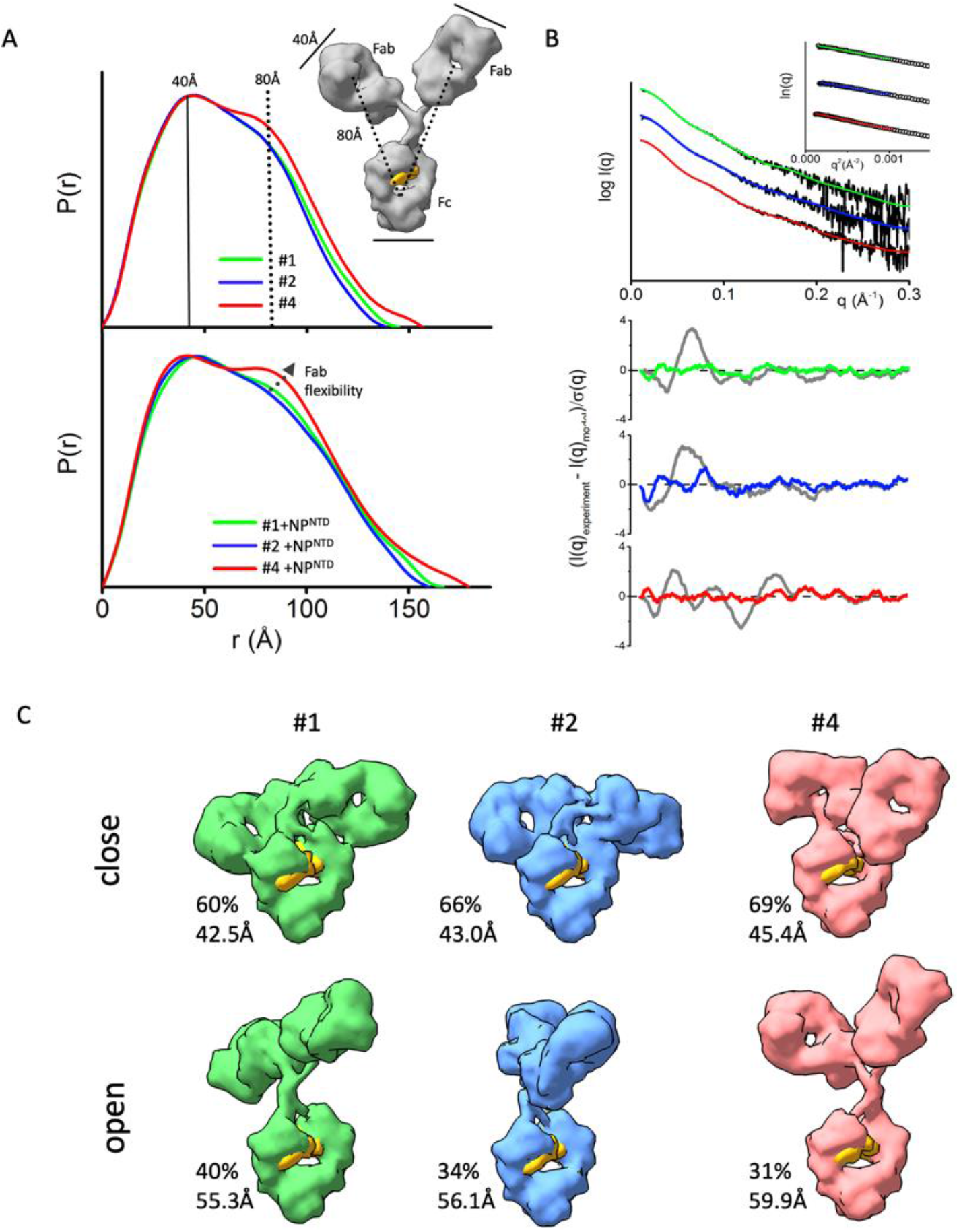
The flexibility of the NP^NTD^-binding mAbs. **A)** P(r) functions for free mAb 1, 2, and 4 (top) and their complexes with the NP^NTD^ normalized on to their maxima. The P(r) shoulder at r∼80Å indicates the Fab-Fc separation described within the atomic model of IgG1 (inset). P(r) peak at 40Å corresponds to the average size across Fc or Fab regions. **B)** Experimental SAXS profiles of free mAb 1, 2, and 4 (black) and theoretical SAXS profiles calculated from their respective two-state atomistic model (green, blue, and red) are shown in the panel. Residuals (Experiment/Model) for the fits of two-state models (green, blue, and red) are shown together with the best single model (gray) and indicate that the two-state model is required to match the experimental SAXS curves. **C**) Two-state models for free mAb 1, 2, and 4 are shown together with the corresponding weights in % and Rg values. The Rg values and weights of the mAb4 further confirm a larger separation between the Fc and Fab region. The atomistic models are shown as molecular envelops at 10Å resolution. The glycan-moiety in the Fc region is colored yellow.

### mAbs with distinct flexibility of the Fab domains

Interpretation of SAXS and P(r) functions is further enhanced by available atomic models of mAbs. While the crystal structure of intact human IgG1 antibody (PDBID:1HZH) does not fit the SAXS data, it forms the basis for creating an ensemble of conformations. We used the program BILBOMD (Pelikan, Hura, & Hammel, 2009) to explore the Fab regions’ conformational space relative to the Fc. BILBOMD performs minimal MD simulations on the Fc-hinge regions at very high temperature, where the additional kinetic energy prevents the Fabs from becoming trapped in a local minimum. This conformational sampling provides a pool of atomistic models (> 10,000) from which SAXS curves are calculated (Schneidman-Duhovny, Hammel, Tainer, & Sali, 2013) and compared to the experimental curve. MultiFoXS algorithm (Schneidman-Duhovny, Hammel, Tainer, & Sali, 2016) is used to identify the weighting of models that fit the experimental data.

At least two distinct conformations are required to fit the SAXS data measured for the three mAbs that bind antigen (mAb 1, 2, and 4). A single conformation from BILBOMD failed to adequately match our measured SAXS profiles (Figure 2B). For each mAb, we found one closed conformer, defined by R_g_ values between 43-45Å, and one open conformer, defined by Rg values between 50-60 Å. These conformations are shown in Figure 2C for each of the mAbs that bind antigen.

MAb-binders (mAb 1, 2, and 4) show differences in conformational variability between two-states. Both mAb4 open and closed conformers show significant separation between Fc and Fab regions (Figure 2C) relative to those found to fit data from the other two mAbs. This difference provides further insight into the prominent P(r) shoulder observed for mAb4 (Figure 2A).

The same feature, indicating additional mAb4 flexibility, is observed in the P(r) functions when NP^NTD^ is present (Figure 2A). A more distinct separation of the P(r) shoulder in the mAb4-NP^NTD^ complex and free state (Figure 2A bottom) indicates larger distancing of Fab from Fc. On the other hand, small P(r) shoulders (Figure 2A) together with small experimental Rg values (Figure 1B) of the mAb1-NP^NTD^ and mAb2-NP^NTD^ complexes correlate with the P(r) shapes of free mAb1 and mAb2, which suggests rigidity of the antibodies. Comparable Fab-flexibility between free and NP^NTD^-bound states agree with previous MD simulations showing only minor allosteric communication between Fab and Fc domains upon antigen binding (Zhao, Nussinov, & Ma, 2019).

### Fab flexibility correlates with sandwich or linear pairing of mAbs

MAb pairs that simultaneously bind the same NP^NTD^ through different epitopes are also readily distinguished from pairs that compete for the same epitope by SEC-MALS-SAXS. Based on the SEC elution profile and MALS-determined molecular weight across the SEC peak, we show that the NP^NTD^ does not bridge mAb1 and 4 (Figure 3A). Thus, mAb1 and 4 compete for binding to NP^NTD^. In contrast, higher mass species were formed by mixing mAb2-NP^NTD^ with either mAb1 or mAb 4, showing that mAb1-2 or mAb2-4 are pairing through simultaneous binding with NP^NTD^ at different epitopes (Figure. 3A). Control experiments show that neither mAb1-2 nor mAb2-4 mixtures form larger complexes in the absence of NP^NTD^ (Supplementary Figure 2).

**Figure 3.**
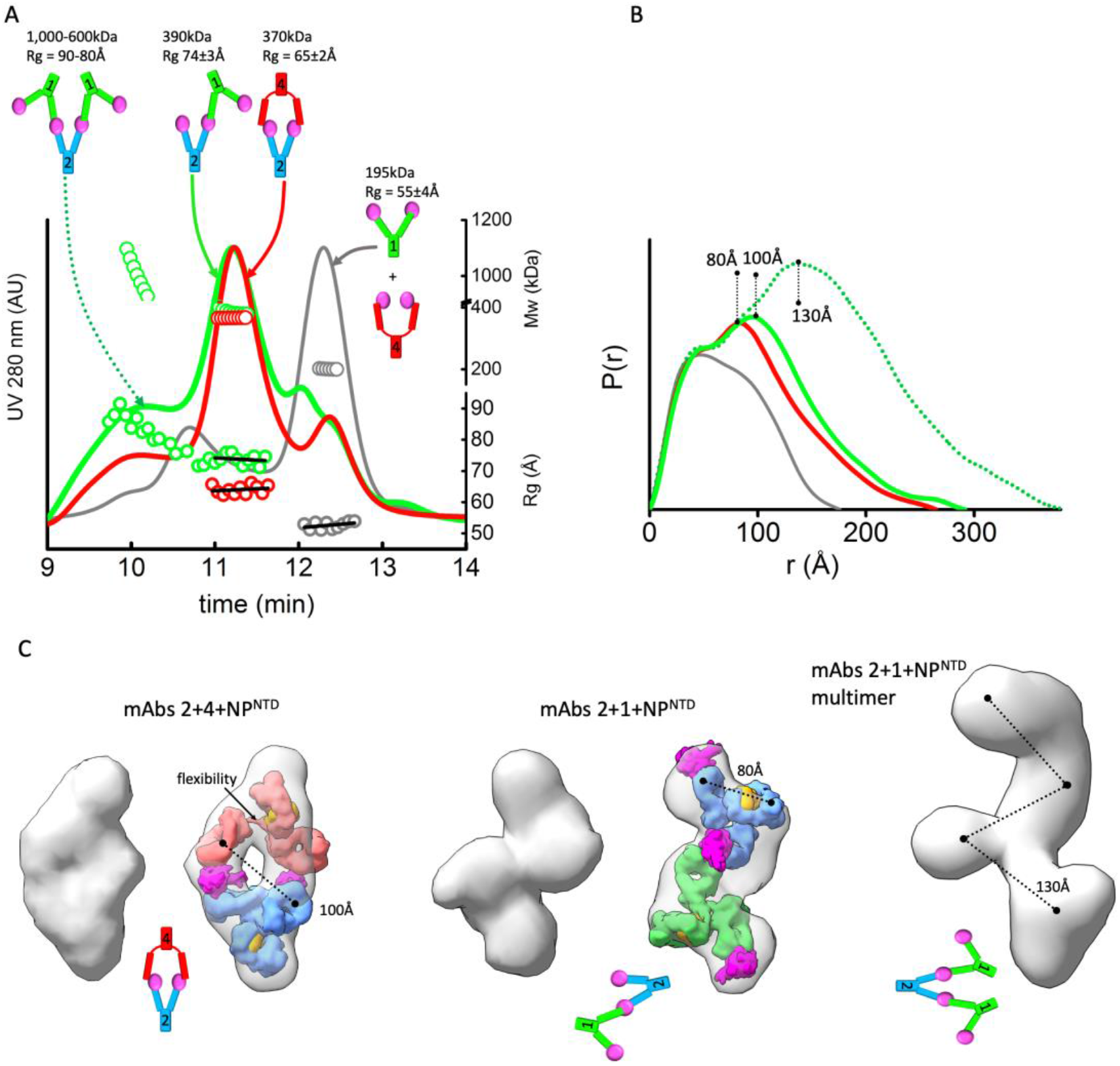
MAbs linear or sandwich pairing depends on inherent flexibility. **A)** SEC-MALS-SAXS chromatograms for the mAb1-2-NP^NTD^ (green), mAb2-4-NP^NTD^ (red) and mAb1-4-NP^NTD^ (gray) samples. Solid lines represent the UV 280nm signal in arbitrary units, while symbols represent molecular mass (top) calculated from MALS and Rg values (bottom) for each collected SAXS frame versus elution time. **B)** P(r) functions calculated for the experimental SAXS curves for the main SEC peak of mAb1-2-NP^NTD^ (green), mAb2-4-NP^NTD^ (red), mAb1-4-NP^NTD^ (gray), and early SEC shoulder of mAb1-2-NP^NTD^ (green dots). The P(r) functions are normalized at the r=40Å. The P(r)-maxima peaks are indicated. Experimental SAXS and Guinier plots are shown in Supplemental Figure 1. **C)** Average SAXS envelope obtained for mAb2-4-NP^NTD^, mAb1-2-NP^NTD^ complex were calculated using P2 symmetry operator. Average SAXS envelopes calculated using P1 symmetry operator are shown in Supplemental Figure 3. A single representative envelope was manually superimposed with compact conformers of mAb1 (red), mAb2 (blue), and mAb4 (green) taken from the two-state model of free mAbs (see Figure 2C). The structure of NP^NTD^ (magenta; PDB ID: 6VYO) was manually docked at the proximity of the CRD3 -Fab region. Additionally, the SAXS envelope obtained for the larger multimer of mAb1-2-NP^NTD^ determined in P1 symmetry is shown.

Each mAb pair binds NP^NTD^ in different stoichiometries and orientations. Mass by MALS and SAXS from the main elution peak show the complex formed by mAbs1-2 is ∼380 kDa, while the mAbs2-4 is ∼340 kDa; corresponding to two antibodies bound by three or two NP^NTD^ molecules, respectively. Also, the orientation of binding between the pairs is very different. The Rg of mAb1-2-NP^NTD^ is 74 Å relative to the 68 Å measured for mAb2-4-NP^NTD^ (Figure 3A, bottom right axis). Furthermore, Rg changes are accompanied by a shift in the secondary peak in the P(r) distribution (100Å vs. 80Å). To gain insights into the structures these mAb pairs form, we reconstructed SAXS envelopes for both (mAb1-2-NP^NTD^, mAb2-4-NP^NTD^). The envelopes for mAb2-4-NP^NTD^ show a sandwich-like assembly with a hollow feature in the center of the model, whereas the mAb1-2-NP^NTD^ adopts a linear arrangement.

We manually superimposed the SAXS envelopes with their corresponding mAb-atomistic models to approximate the overall arrangement of mAb-pairs. The sandwich-like arrangement of mAb2-4-NP^NTD^ matches the SAXS envelope and shows two antigens bound between two Fabs. The SAXS envelop of mAb1-2-NP^NTD^ matches a linear arrangement of the antibodies where only one NP^NTD^ is shared between mAb1-2 (Figure 3B). The shapes and models of the complex provide insights into the P(r) distributions’ shifts.

We postulate that the difference in orientation fundamentally relies on differences in the flexibility of the mAbs. The mAb2-4 pair contains the flexible mAb4 and shows a closed and capped arrangement around two antigens. MAb4’s flexibility allows the Fab-regions stretch to accommodate two NP^NTD^ molecules’ binding located on the Fabs of mAb2. In contrast, the more rigid mAb1-Fab regions prohibit the Fabs’ positioning onto the NP^NTD^ located on the mAb2. Thus, the relative rigidity of both mAb2 and mAb1 enforces the linear arrangement of the mAb1-2-NP^NTD^ complex.

The linear antibody-antigen arrangement of the mAb1-2-NP^NTD^ complex should permit further networking of multiple mAbs through the uncovered epitopes of the NP^NTD^ molecules bound to the outermost Fab regions. Indeed, there is a notable presence of very large complexes (∼1MDa) in the mAb1-2-NP^NTD^ sample (Figure 3A), suggesting further elongation of the complex by extending the rigid linear arrangement (Figure 3BC). These observations suggest flexibility of mAbs is a factor in the agglutination of mAb - antigen complexes.

### SPR kinetic analysis revealed comparable picomolar affinities of all antibodies

To compare the relative affinity of each mAb for antigen, we performed binding kinetic assays. In addition, we performed assays on HRP-conjugated mAbs in preparation for ELISAs, described below. Due to the high affinities of the mAbs, we opted to use a kinetic titration (single cycle kinetics) strategy and avoid problematic regeneration steps (Methods). We measured the binding kinetics of mAb 1, 1-HRP, 2, 4, and 4-HRP by surface plasmon resonance (SPR) (Supplementary Figure 4). All antibodies (unconjugated and HRP-conjugated) had high-affinity constants (K_D_) in the picomolar range (Supplementary Table 1). The K_D_ of HRP-conjugated mAb1 and 4 are very similar, at 11 and 28 pM, respectively. The percent activity of the HRP-conjugated antibodies is lower than unconjugated, suggesting that conjugating HRP on the antibodies affects the percentage of available antibodies for interaction on the SPR sensor chip. The possibility exists that this effect could also be present in the chip-free solution-based ELISA assay. However, the high concentration of HRP-conjugated antibodies used (0.4 mg/mL; Methods), relative to the picomolar affinities, represents a large excess of functional, high-affinity HRP-conjugated antibodies in the ELISA assay. Therefore, the antibodies have comparable kinetics, effectively excluding them as explanations for functional outcomes.

### A modified ELISA protocol “boosts” the signal of the linear mAb arrangement

We sought to assess the consequences of the observed mAb linear arrangement vs. sandwich pairing (Figure 3) on detection limits. Since mAb2 pairs with mAb1 and mAb4, we used mAb2 as the NP^NTD^ capture antibody and conjugated HRP to mAb1 and mAb4 (1-HRP, 4-HRP) to serve as the detection antibodies. We hypothesized that the linear arrangement of mAb1-2-NP^NTD^ could facilitate two or more 1-HRP antibodies to capture mAb2 on the plate, leading to a boost in the signal. This would contrast with the sandwich pairing of mAb4, which closes off the further binding and constrains assembly to a 1:1 ratio of 4-HRP to mAb2. To test this hypothesis, we developed a modified ELISA protocol. To enhance detection, we modified the standard ELISA protocol. The main two differences between this and a standard ELISA are: 1) The detection HRP-conjugated mAbs are added directly on top of the samples during the incubation period that is typically used for the capture of the antigen only, and 2) Free (non-plate-bound) mAb2 is “spiked” into the detection HRP-conjugated mAb solutions before their addition on top of the samples. We rationalized that adding antigen simultaneously with detection antibodies would initiate maximal polymerization and that a later “spiking” in of mAb2 would further extend “networking” of the linear mAb2-1-NP^NTD^ arrangement (Figure 3 – middle/right panels). Whereas, sandwich pairing of mAb2-4-NP^NTD^ does not allow the polymerization of antibodies (Figure 3 – left panel).

Following this protocol, we observe improvements in detection limits using mAb2-1 - NP^NTD^ relative to mAb2-4-NP^NTD^ (Figure 4). In repeated experiments, Figure 4AB, the signal of 1-HRP is ∼2-fold higher than 4-HRP, although the magnitude of the effect is diminished with decreasing concentration of antigen (Figure 4AB - insert; Figure 4C). The two independent experiments (Figure 4AB) had a control experiment done in parallel on the same plate. The control experiment follows the standard ELISA protocol, where the plate was washed prior to the addition of the mAb-HRP for a twenty-minute incubation (Supplementary Figure 5A). To control for the longer incubation time of the mAb-HRP with the samples of our modified ELISA protocol, we ran an additional control (Supplementary Figure 5B), where the mAb-HRP had a longer incubation time of 1.5 hrs. No “boost” of the 1-HRP signal over the 4-HRP level was seen in either control experiment (Supplementary Figure 5AB). Simultaneously, there was a general elevation of both signals in the one with the longer incubation time (Supplementary Figure 5B). These results collectively demonstrate the ability to capitalize on the linear-mAb arrangement functionally, which results from the structural rigidity of the antibodies (Figures 1-3).

**Figure 4.**
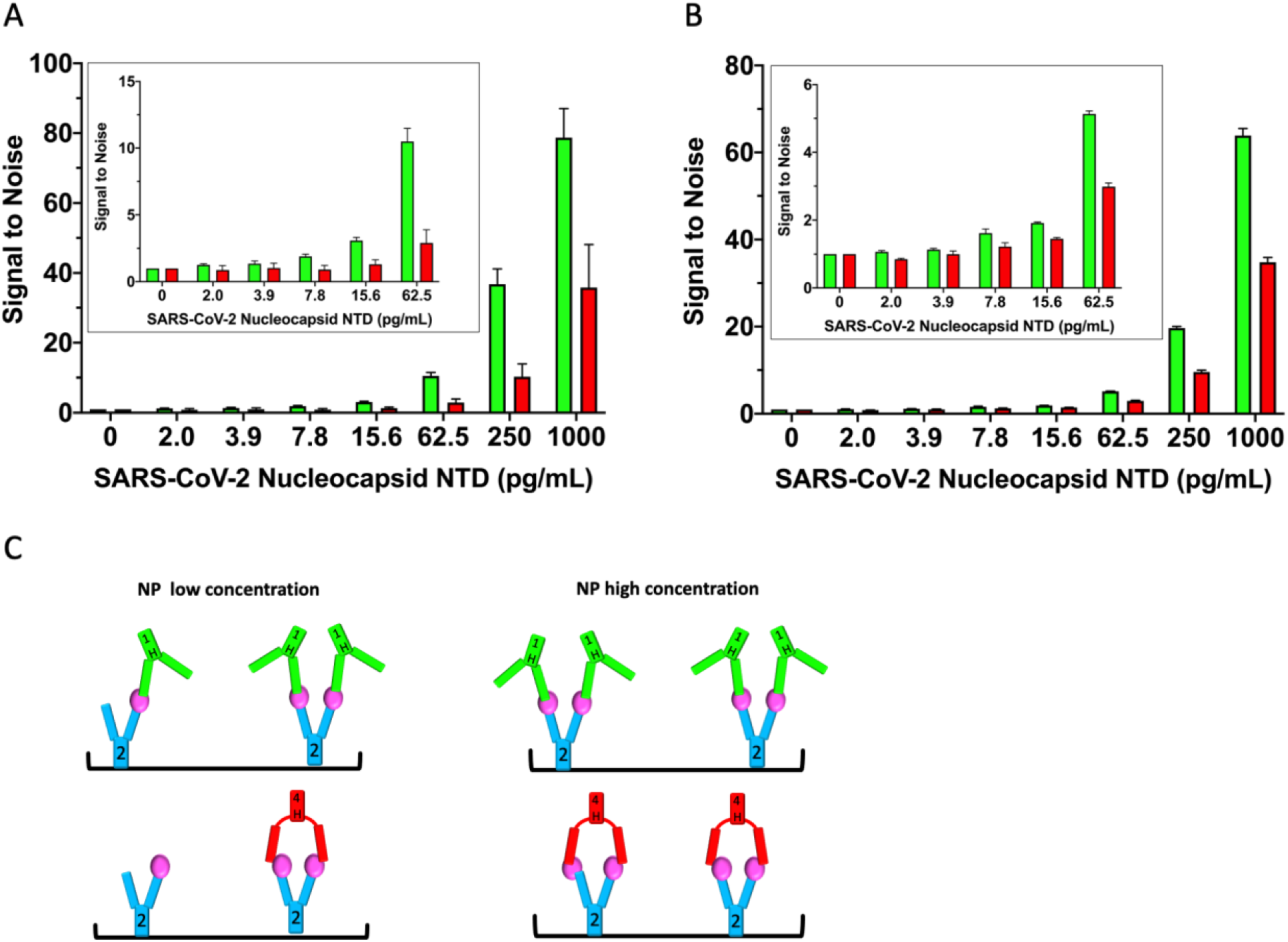
Linearly arranged mAbs show boosted signal in modified ELISA. **A)** A modified ELISA where the detection HRP-conjugated mAbs (1-HRP in green, 4-HRP in red) are added directly on top of the samples during the NP^NTD^ capture incubation period. Free (non-plate-bound) mAb2 is “spiked” into the detection HRP-conjugated mAb solutions before their addition on top of the samples. The corresponding standard control ELISA protocol run in parallel on the same plate is shown in Supplementary Figure 5A. **B)** Repeat of the experiment conducted in A, with a corresponding control ELISA protocol run in parallel on the same plate with a longer mAb-HRP-sample incubation period, shown in Supplementary Figure 5B. **C)** Schematic of low versus high concentration of NP^NTD^ in samples. In both experiments, the 1-HRP that forms the more rigid linear arrangement in the unconjugated form (mAb2-1-NP^NTD^) shows an ∼2-fold increased ELISA signal, relative to 4-HRP, that forms a sandwich arrangement in the unconjugated form (mAb2-4-NP^NTD^).

Further, we were interested in whether the structurally enforced functional “boost” effect could be maintained in the presence of a virion-disrupting detergent (Supplementary Figure 6) since NP is present inside virions. SARS-CoV-2 virions are not lysed adequately in the presence of 0.5% Tween-20, a commonly used detergent in ELISAs that is present in our protocol at a lower concentration, 0.05% (Methods), but were effectively lysed in the presence of 0.5% triton X-100 (Patterson et al., 2020). Therefore, we used our same modified ELISA protocol that demonstrated the “boost” (Figure 4AB), except that we used PBS pH 7.4 plus 0.5% triton X-100 as the sample dilution buffer, instead of PBS pH 7.4 alone. The presence of triton X-100 reduced the “boost”, although it is still detectable (Supplementary Figure 6). Interestingly, the presence of triton X-100 appears to have increased the overall limit of detection (LOD) to lower than 0.4 pg/mL. In contrast, it is clearly not this low in the detergent’s absence (compare 0 and 2 pg/mL in Figure 4AB with 0 and 1.7 pg/mL in Supplementary Figure 6). Further improvements could be gained to maximize both the “boost” and detergent effects. Together, these results suggest that combining our modified ELISA protocol with the presence of a SARS-CoV-2 virion lysing concentration of triton X-100 leads to a highly sensitive ELISA assay, with great potential for further diagnostic development.

## Discussion

The ongoing SARS-CoV-2 pandemic has highlighted the need for sensitive point-of-care diagnostics (POCs), which are primarily antibody-based technologies (Alpdagtas et al., 2020). Currently, mAbs are widely used to detect antigen molecules, including the nucleocapsid protein from SARS-CoV-2. The US Food and Drug Administration recently authorized a lateral flow antigen test as the first over-the-counter, fully at-home diagnostic test for the qualitative detection of SARS-CoV-2 nucleocapsid antigens (FDA, 2020). However, there is an urgent need to improve the detection limit of these diagnostic devices that use various types of colorimetric mAb-based assays (Alpdagtas et al., 2020). Multiple approaches, like florescent immunoassays (Ahn et al., 2009; Loeffelholz & Tang, 2020; Wang et al., 2020), nanoparticle luminescence (Hesari & Ding, 2020; Teengam et al., 2017) or magnetic beads as the antibody support surface (Fabiani et al., 2021), are used to enhance the detection of antigen. In immunoassays and RT-PCR, the detection is signaled through chemical conjugation to an enzyme or nanoparticle that drives a colorimetric reaction, a fluorophore, or another moiety. However, a limitation of existing immunoassays in detecting antigens relative to RT-PCR is the lack of exponential amplification of signal when probes detect an antigen. Immunoassays mainly rely on antibody-antigen binding at a 1:1 ratio. An immunoassay diagnostic with a greater detection-to-capture antibody ratio will also have a greater signal-to-antigen ratio, effectively enhancing overall specific antigen detection.

Despite the widespread use of antibodies in diagnostics and treatments, an understanding of structural properties that affect antibody-antigen interactions in their aqueous environment is currently insufficient to guide optimization for these purposes. Here, we describe how antibodies’ inherent flexibility leads to distinct antigen-binding arrangements that drive different functional outcomes in a SARS-CoV-2 detection ELISA. We show that we can rapidly assess new antibody-antigen interactions using SEC-MALS-SAXS to identify pairs that bind in a linear arrangement (Figure 3), which results in a more sensitive detection assay (Figure 4). A critical benefit of using the SEC-MALS-SAXS approach is its ease of use and the ability to study antibody-interactions in solution. It has previously been shown that other techniques that rely on grids (EM) or crystals often do not reflect the dynamic nature of antibodies in solution (Jay et al., 2018).

A central finding in this work is the observation and rationalization of how mAb flexibility can impact larger assemblies of more than one mAb with or without antigen. Many mAbs are abandoned as formulations despite a high affinity for their antigen because of a propensity to aggregate. SEC-MALS-SAXS is a rapid method that allows separation of larger-scale assemblies from single mAbs and, therefore, an interrogation of aggregation propensity on the structural properties of the underlying single mAb. Further, SAXS provides the resolution to distinguish rigid and flexible mAbs in solution. Flexibility can arise not only from primary sequence differences but also due to glycosylation. This may have been a factor in our studies as we used both mouse (mAb1, mAb2) and rabbit (mAb4) host antibodies (Yagi, Yanaka, & Kato, 2018). In light of our results on the impact of flexibility, further studies to assess glycosylation’s effect may be worthwhile as glycosylation may be adjusted.

Having identified two mAbs (mAb1 and mAb4) with differing degrees of flexibility (Figure 2) that can both bind SARS-CoV-2 NP^NTD^ simultaneously with a third antibody mAb2, we were able to contrast the larger assemblies composed of mAb2-1 NP^NTD^ versus mAb2-4 NP^NTD^ (Figure 3). Further analysis confirmed the existence of two different binding modes of antibody-antigen-antibody, sandwich, and linear (Figure 3). The linear mode suggested further polymerization might be possible. This polymerization would be considered aggregation when interpreted by other methods. However, we sought to use this propensity to overcome the limitation on amplification inherent in antigen-based diagnostics.

Based on the above observations, we developed a modified ELISA. We used mAb2 as a common capture antibody and mAbs 1 and 4 as detection antibodies. We saw the increased signal for the mAb2-1 pairing, relative to mAb2-4, where mAb1 and mAb4 were coupled to horseradish peroxidase (HRP); a common signal-generating enzyme used in ELISA assays (Lin, 2015). To control for changes in binding kinetics, we used SPR to demonstrate that the near-equivalent binding affinities for nucleocapsid are maintained post-HRP conjugation between the two antibodies. The modified ELISA that employed the linearly arranged mAb2-1 pair consistently generated a larger signal than the sandwich mAb2-4 pair. Further, the signal of 1-HRP is ∼2-fold higher than 4-HRP, and the effect is diminished with decreasing concentration of antigen, which supports our hypothesis.

Having made gains in detection by considering the structural properties of mAbs, more optimization is likely possible. Introducing further rigidity in mAb2 through glycosylation modifications, binding factors like protein A or G, detergents, or other metabolites could enhance further networking and, therefore, detection. The positive signal line in LFAs is often generated by antibodies conjugated to colloidal gold or latex, which accumulate into pink or blue lines, respectively (Tang et al., 2009). These are meant for visual inspection by non-experts in POC devices. A previous study demonstrated that the detection limit of an LFA could be lowered 3-fold, from 3.1 ng/mL to 0.9 ng/mL for detection of aflatoxin B2 in food, through non-covalently clustering (16nm diameter) gold nanoparticles for a visual readout (Tang et al., 2009). Thus, the clustering of signal molecules coupled to antigen-specific antibodies is a viable strategy for lowering the LOD in LFAs. Our study shows that this can be achieved without introducing an additional factor, instead by taking advantage of the antibodies’ structural rigidity. A survey of commercial ELISAs suggests a common LOD of 100 pg/mL, with the most sensitive being 0.01 pg/mL for protein analytes (S. Zhang, Garcia-D’Angeli, Brennan, & Huo, 2014). Many antibodies have been identified against NP and other antigen targets from SARS-CoV-2. SEC-MALS-SAXS could be applied to hundreds in a short amount of time to identify the most rigid. By combining this novel strategy with other optimization methods (e.g., tuning antibody affinities, selecting signaling molecule/moiety), the LOD of standard LFA POC devices could achieve as yet unattained sensitivity for current and future pathogens.

### Materials and Methods Monoclonal Antibody Sources

Seven antibodies were purchased from SinoBiological, and two antibodies from CreativeBiolabs. The catalog numbers for the SinoBiological antibodies are as follows: #1: 40143-MM05, #2: 40143-MM08, #3: 40143-R001, #4: 40143-R004, #5: 40143-R019, #6: 40143-R040, #15: 40588-R0004. The catalog numbers for the CreativeBiolabs antibodies are as follows: #18: MRO-0015YJ, #19: MRO-0016YJ. Antibodies #1 and #4 were chemically conjugated to HRP by SinoBiological CRO Services. The molar HRP:Ab ratio was 2.81 for #1-HRP, and 3.5 for #4-HRP.

### Expression and purification of NP^NTD^

Gene fragment coding of nucleocapsid protein from SARS-CoV-2 was codon-optimized for efficient expression in *E. coli*. The coding sequence of NP^NTD^ comprising residues Asn47 to Ala173 (UniProtKB - P0DTC9) was synthesized and cloned into pMCSG53 vector (Eschenfeldt et al., 2013) by Twist Biosciences, USA. Cloning into pMCSG53 vector introduced to NP^NTD^, a His6-Tag at the N-terminus followed by a cleavage site for tobacco etch virus (TEV) protease. For NP^NTD^ expression, the plasmid was transformed into *E. coli* BL21(DE3)-Gold cells (Strategene) using heat-shock. After transformation, bacterial cells were precultured overnight at 37°C in 100 ml of LB Lennox medium supplemented with 40 mM K2HPO_4_ and 160 mg/L of ampicillin. Subsequently, 40 ml of overnight cultures were used to inoculate 4 liters of LB with 40 mM K_2_HPO_4_ and 160 mg/L ampicillin. Next, cells were incubated at 37°C with 180 RPM shaking for approximately 3 hours until reaching optical density at 600 nm equal to 1. Subsequently, the bacteria culture was cooled down for 1 hour in an incubator set to 4°C with 180 RPM shaking. Expression of NP^NTD^ was induced with 0.2 mM isopropyl β-d-1-thiogalactopyranoside, supplemented with 0.1% glucose, and incubated overnight at 16°C. Bacteria cells were harvested by centrifugation at 4°C, 5000 RCF for 10 minutes. Cell pellets were resuspended in lysis buffer 50 mM HEPES pH 8.0, 500 mM NaCl, 5% v/v glycerol, 20 mM imidazole, and 10 mM β-mercaptoethanol (1 g of cells: 5 ml of lysis buffer) for purification or frozen and stored at −80°C until purification.

After overexpression of NP^NTD^, bacteria cells were lysed by sonication on ice using 120W output power for 5 minutes (4 sec pulses of sonication followed by 20 sec brakes). After sonication, samples were centrifuged to remove cellular debris (30k RCF, 4°C, 1 hour). We used a vacuum-assisted purification system to perform NP^NTD^ purification with immobilized metal affinity chromatography (IMAC). Using 5 ml of Ni^2+^ Sepharose (GE Healthcare) loaded on a Flex-Column (420400-2510) attached to a Vac-Man vacuum system (Promega), beads were equilibrated in a lysis buffer. The cell lysate was loaded on the column, and Ni^2+^ Sepharose was washed using 20 column volumes of lysis buffer. For elution, lysis buffer was supplemented with imidazole up to 500 mM (pH 8). After elution for removing 6His-Tag, we used TEV protease added in a molar ratio 1 TEV to 40 NP^NTD^. TEV cleavage leaves 3 residues SerAsnAla at the N-terminus of NP^NTD^. Next, NP^NTD^ was concentrated using 10kDa cut-off centrifugal protein concentrators (Merck-Millipore). Subsequently, we performed SEC of NP^NTD^ using a Superdex S200 16/600 column attached to an Åkta Express (GE Healthcare) purification system. SEC was done at 4°C in a buffer containing 20 mM HEPES, 500 mM NaCl, 5% v/v glycerol, 10 mM β-mercaptoethanol, pH 8.0. Purified fractions of NP^NTD^ from the middle of the gel filtration elution peak were concentrated to 10.7 mg/ml. Protein was flash cooled using 40 μl aliquots dropped directly into liquid nitrogen. Samples were stored at −80°C or on dry ice during shipment. Upon thawing, samples were stored at 4°C and diluted in PBS pH 7.4 for functional assays.

### Small-angle X-ray scattering and Multi-Angle Light Scattering data acquisition in line with SEC (SEC-MALS-SAXS)

For SEC-MALS-SAXS experiments, 60 μL of samples containing mAb ∼1-3 mg/mL and NP^NTD^ in 1:5 molar ratio were prepared in PBS pH 7.4 buffer. mAb 1, 2, and 4 were also measured in the absence of NP^NTD^ using the same buffer conditions. The mAb pairs 1-2, 1-4, and 2-4 in the presence of NP^NTD^ were prepared in the molar ratio of 1:1:10 in the same buffer conditions. All samples were incubated for a minimum of 30 minutes before the injection on SEC.

SEC-MALS-SAXS were collected at the ALS beamline 12.3.1 (Dyer et al., 2014). X-ray wavelength was set at λ=1.127 Å, and the sample to detector distance was 2100 mm, resulting in scattering vectors, q, ranging from 0.01 Å^-1^ to 0.4 Å^-1^. The scattering vector is defined as q = 4πsinθ/λ, where 2θ is the scattering angle. All experiments were performed at 20°C, and data were processed as described (Hura et al., 2009). Briefly, a SAXS flow cell was directly coupled with an online Agilent 1260 Infinity HPLC system using a Shodex KW 803 column. The column was equilibrated with running buffer (PBS pH 7.4) with a 0.5 mL/min flow rate. 55 µL of each sample was run through the SEC, and three second X-ray exposures were collected continuously during a 30 min elution. The SAXS frames recorded before the protein elution peak were used to subtract all other frames. The subtracted frames were investigated by the radius of gyration (Rg) derived by the Guinier approximation I(q) = I(0) exp(-q^2^*Rg*2/3) with the limits q*Rg<1.5. The elution peak was mapped by comparing integral ratios to background and Rg relative to the recorded frame using the program SCÅTTER. Uniform Rg values across an elution peak represent a homogenous state of mAb or its complex. Final merged SAXS profiles (Supplementary Figure 1), derived by integrating multiple frames across the elution peak, were used for further analysis, including a Guinier plot, which determined the aggregation free state (Supplementary Figure 1). The program SCATTER was used to compute the pair distribution function (P(r)) (Figure 1B and 3C). The distance r where P(r) approaches zero intensity identifies the macromolecule’s maximal dimension (Dmax, Figure 1B, Table 1). P(r) functions for single mAb (Figure 2A), single mAb+NP^NTD^ (Figure 1A) were normalized at the maxima except the P(r) of NP^NTD^ alone (Figure 1B); mAb1-2-NP^NTD^ and mAb2-4-NP^NTD^ complexes (Figure 3B) where the area under P(r) function correlates to molecular weight estimated by SAXS.

The Eluent was subsequently split 3 to 1 between the SAXS line and a series of UV at 280 and 260 nm, MALS, quasi-elastic light scattering (QELS), and refractometer detector. MALS experiments were performed using an 18-angle DAWN HELEOS II light scattering detector connected in tandem to an Optilab refractive index concentration detector (*Wyatt Technology*). System normalization and calibration were performed with bovine serum albumin using a 45 μL sample at 10 mg/mL in the same SEC running buffer and a dn/dc value of 0.19. The light scattering experiments were used to perform analytical scale chromatographic separations for molecular weight (MW) determination of the SEC analysis. UV, MALS, and differential refractive index data were analyzed using Wyatt Astra 7 software to monitor the sample’s homogeneity across the elution peak complementary to the above-mentioned SEC-SAXS signal validation.

### Solution state modeling

BILBOMD (Pelikan et al., 2009) rigid body modeling along with a FoXS and MultiFOXS (Schneidman-Duhovny et al., 2013, 2016) approach was used to define, select and weight the two-state atomistic model that best agree with individual SAXS profiles of free mAb1, 2 and 4. The crystal structure with PDB ID: 1HZH (Saphire et al., 2001), including the glycans moiety, was used as an initial model. In the case of glycoproteins, the glycans’ contribution to the scattering is known to be larger than protein alone (Hammel et al., 2002). Minimal molecular dynamics (MD) simulation applied on the mAb hinge regions explores the Fab domain’s conformational space relative to the Fc-glycan region. The disulfide bonds in the hinge region were kept intact. In the conformational sampling, individual Fabs would move independently of one another. A single best-fit and two-state models were selected for each mAb using MultiFOXS (Schneidman-Duhovny et al., 2013, 2016).

The SAXS envelopes were reconstructed from the experimental data of mAb2-4-NP^NTD^ and mAb1-2-NP^NTD^ complex using the program DAMMIF (Franke & Svergun, 2009). Ten bead models obtained for each SAXS experiment were averaged by DAMAVER (Volkov & Svergun, 2003) to construct the average model representing each reconstruction’s general structural features. Bead models were converted to volumetric SITUS format with the pdb2vol kernel convolution utility (Wriggers, Milligan, & McCammon, 1999). SAXS data and SAXS derived models have been deposited in SIMPLE SAXS database (https://saxs-server.herokuapp.com/), and experimental SAXS parameters are reported in Table 1.

### SPR

Affinity and kinetic data were acquired using a Biacore T200. All antibodies were coupled to a CM5 Biacore sensor chip using amine coupling. MAbs 1, 2, 4, 4-HRP were coupled using 10 mM acetate pH 5.5 at 10 ug/mL, while mAb 1-HRP required 10 mM acetate pH 5.0 at 10 ug/mL. All experiments were run in 10 mM HEPES pH7.4, 150 mM NaCl, 35 mM EDTA, 0.01% surfactant P20. NP^NTD^ analyte injections for the single-cycle kinetic titrations were as follows: 0.074 nM, 0.22 nM, 0.67 nM, 2 nM, 6 nM. The dissociation time was 3600s.

### ELISA Development

ELISAs were developed using the following materials: Corning 96-Well High-Binding Flat-Bottom Microplates from StemCell (Cat. # 38019), and R&D Systems, Stop Solution 2N Sulfuric Acid (Cat. # DY994), Substrate Reagent Pack (Cat. # DY999), Bovine Serum Albumin-ELISA grade (5217/100G). All reagents were allowed to warm to room temperature before use. ELISA signals were recorded using a POLARstar Omega plate reader. Samples were diluted in PBS unless otherwise stated. The wash buffer consisted of PBS pH 7.4 and 0.05% Tween 20. The blocking buffer consisted of PBS pH 7.4, 2% BSA, and 0.1% Tween 20. Unless specified, the following modified ELISA protocol was used: 100 uL per well of 4 mg/mL mAb2 capture antibody in PBS was added to the ELISA microtiter plate. The plate was sealed and incubated overnight at room temperature. The next day, the solution was discarded and washed with wash buffer three times. The plate was blocked with 300 uL per well of blocking buffer for one hour at room temperature. The plate was washed three times. NP^NTD^ was diluted into PBS as a serial dilution concentration series, and 100 uLs per well was added to the plate. For detection and “antibody networking” assessment, 100 uL of PBS containing 0.4 mg/mL antibody-HRP with or without 0.4 mg/mL mAb2 capture antibody was added directly to the samples in the plate, for a final concentration of 0.2 mg/mL. The plate frame was gently tapped for one minute to mix, sealed, and incubated protected from light for 1.5 hours at room temperature. The plate was washed three times. Before use, the Substrate Reagent was prepared by combining equal parts of component A & B. 100 uL of working Substrate Reagent was added to each well and incubated for 20 minutes at room temperature protected from light. 50 uL of Stop Solution was added to each well, and the plate was gently tapped to ensure thorough mixing. The optical density (OD) of each well was determined within 30 minutes of stopping the reaction. The OD 450nm and 540nm were recorded. The data was background corrected in excel by subtracting the OD 540nm from the 450nm signal. For normalized data, all signals were individually divided by the background signal. All samples were run in triplicate. The mean and standard error of the mean was calculated and plotted in GraphPad Prism.

## Acknowledgements

Funding for this research was provided by federal funds from the DOE through the National Virtual Biotechnology Laboratory, a consortium of DOE national laboratories focused on response to COVID-19, with funding provided by the CARES Act and in part by the National Institute of Allergy and Infectious Diseases, National Institutes of Health, Department of Health and Human Services, under Contract HHSN272201700060C. Funding for the SIBYLS beamline at the Advanced Light Source was provided in part by the Offices of Science and Biological and Environmental Research, U.S. Department of Energy through DOE BER Integrated Diffraction Analysis Technologies (IDAT) program and National Institutes of Health (NIH) grants P01 CA092584). We would like to thank Robert Jedrzejczak and Lukas L. Welk for their help in protein expression and purification. We would also like to thank Andrew Barile-Hill from Cytiva for SPR data collection/analyses.

