## Supplementary Figures 1-6 & Table 1 for "Rigid monoclonal antibodies improve detection of SARS-CoV-2 nucleocapsid protein"

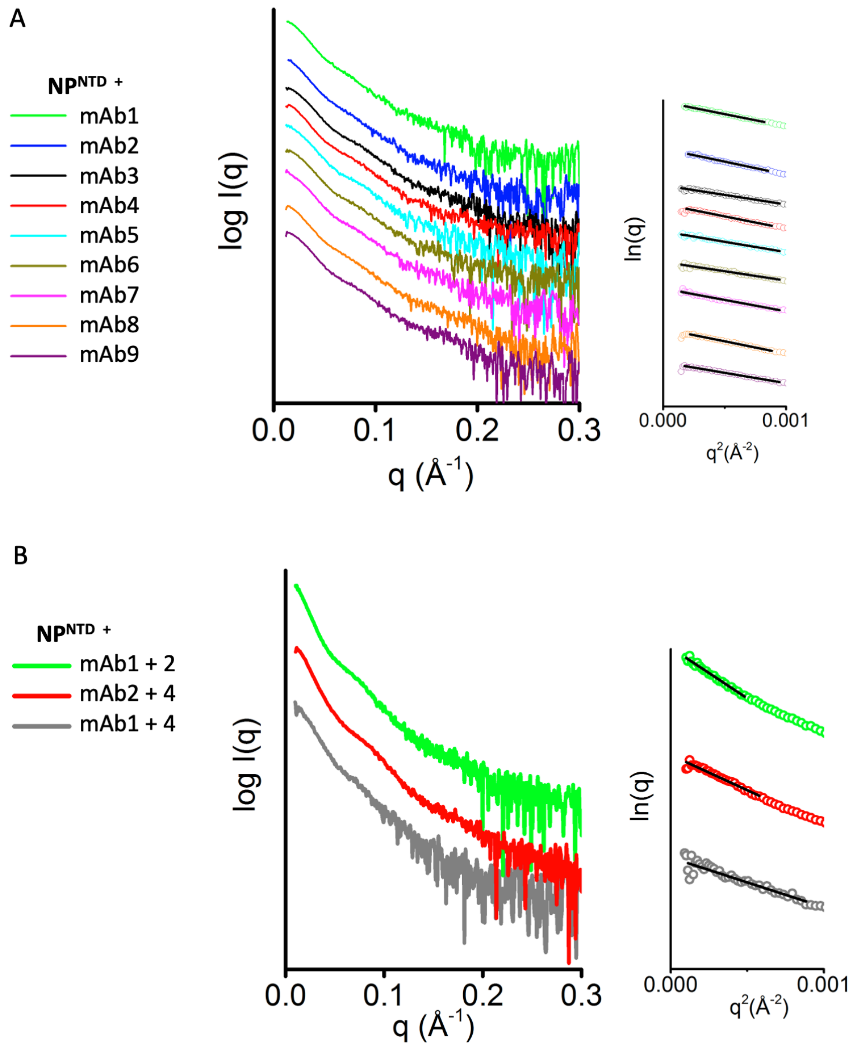


**Supplementary Figure 1. SAXS profiles.**
(A) Experimental SAXS curves of all tested mAb + $NP^{NTD}$ samples (colored as indicated). Right panel: Corresponding Guinier plot with q*Rg <1.5 limit. (B) Experimental SAXS curves for the main SEC peak of mAb1-2-$NP^{NTD}$ (green), mAb2-4-$NP^{NTD}$ (red), mAb1-4-$NP^{NTD}$ (gray). (A-B) Guinier plots were used to determine Rg values used in Figure 1B-inset and listed in Table 1.

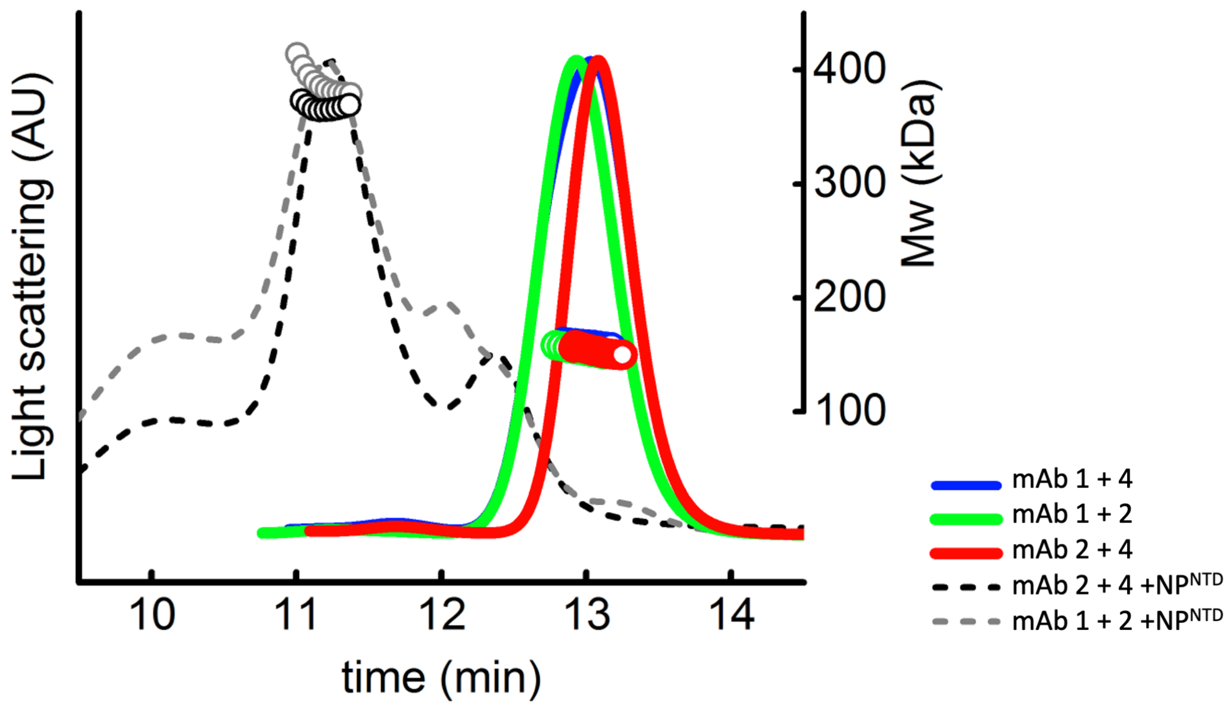


**Supplementary Figure 2. mAb pairs do not form a large assembly in absence of NP$^{NTD}$.**

SEC elution profile (lines) and MALS-determined molecular weight (circles) across the SEC peak for mAb1+4, mAb1+2, mAb2+4 in absence of NP$^{NTD}$ (in comparison to the mAb1+2, mAb2+4 in presence of NP$^{NTD}$ (colored as indicated).

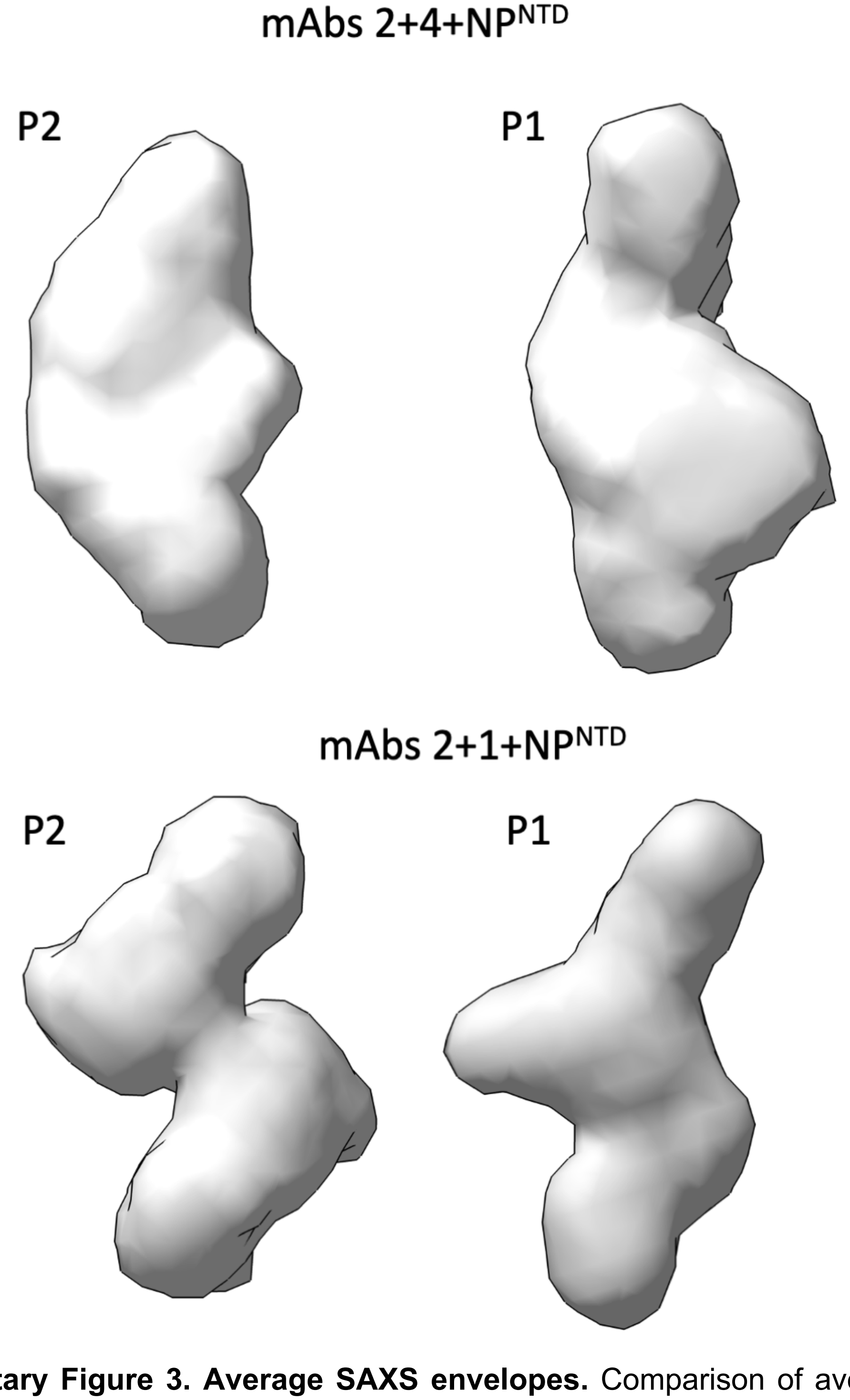


**Supplementary Figure 3. Average SAXS envelopes.** Comparison of average SAXS envelopes obtained for mAb2-4-NP$^{NTD}$, and mAb1-2-NP$^{NTD}$ complexes reconstructed using a P2 (left) and P1 (right) symmetry operator.

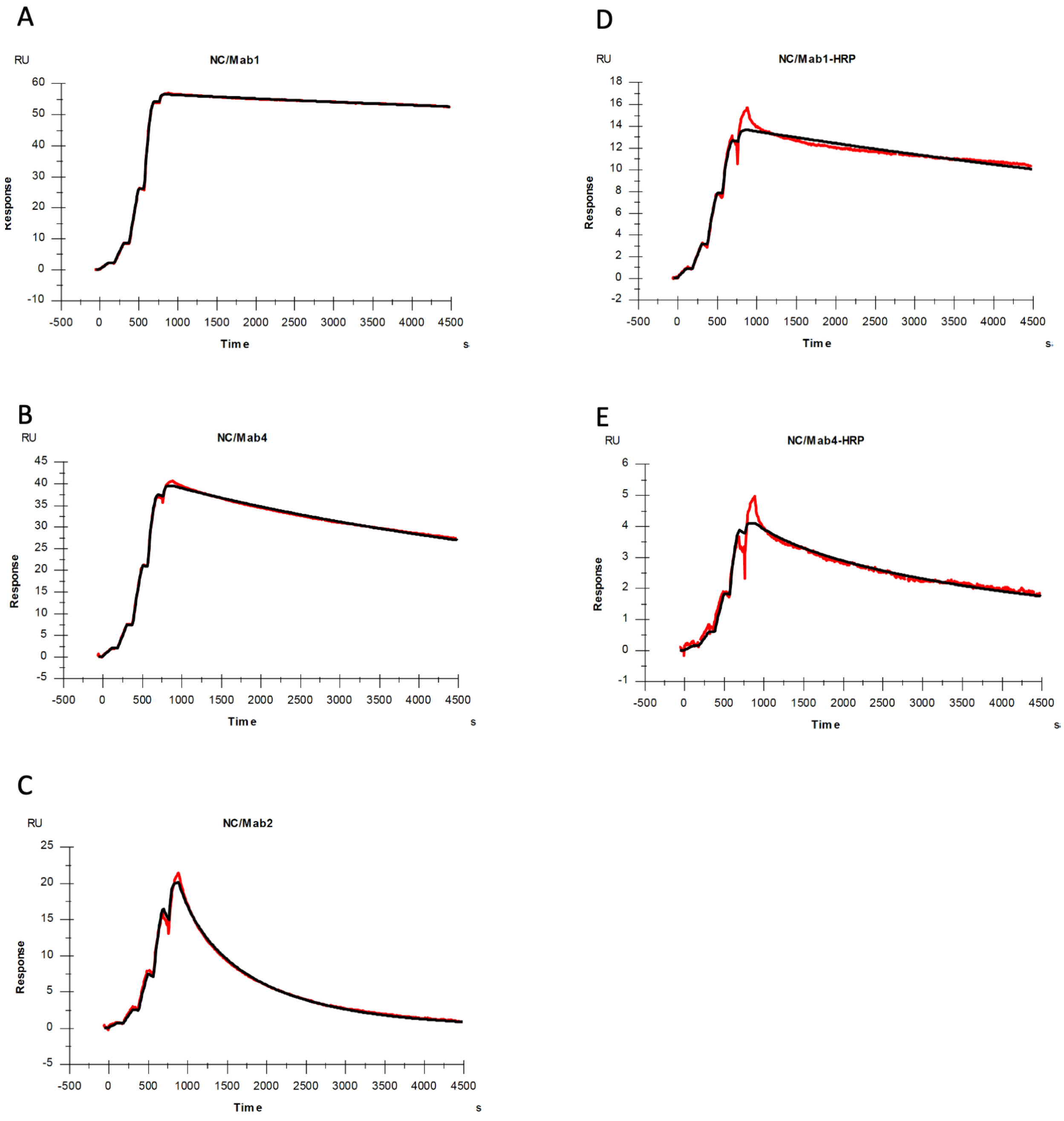


**Supplementary Figure 4. Binding kinetics of $NP^{NTD}$ with the mAb 1, 1-HRP, 2, 4, and 4-HRP by surface plasmon resonance (SPR).**

$NP^{NTD}$ at five concentrations (0.074nM, .22nM, .67nM, 2nM, 6nM ) were tested using single-cycle kinetics with immobilized , mAb1 (A), mAb4 (B), mAb2 (C), mAb1-HRP (D), and mAb4-HRP (E).

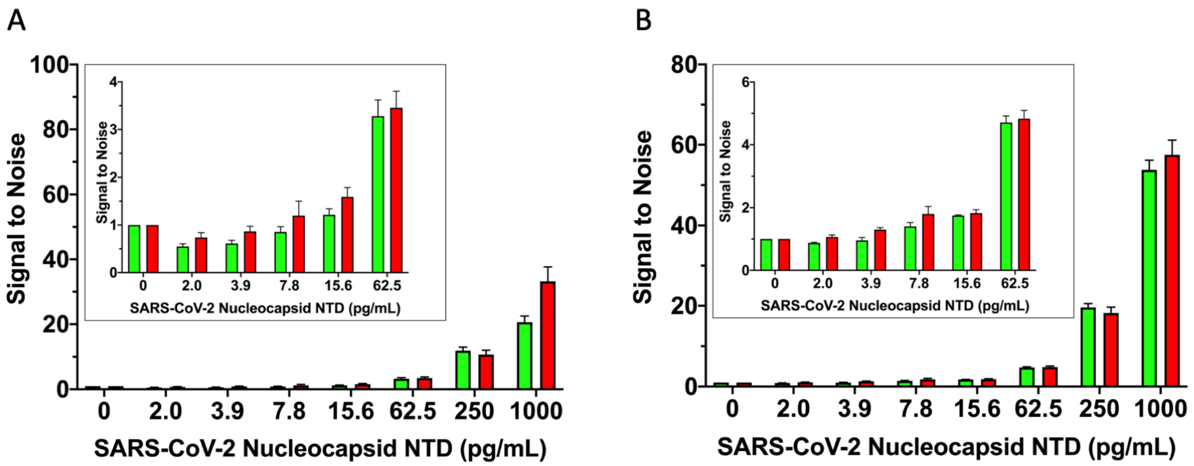


**Supplementary Figure 5. Standard ELISA control protocol shows no boost.**
(A) A standard ELISA where the plate was washed prior to addition of the detection HRP-conjugated mAbs (1-HRP in green, 4-HRP in red), for a twenty minute incubation. (B) A repeat of the experiment shown in A, except the mAb-HRPs had a longer incubation time of 1.5 hrs. In both experiments, no boost is observed, regardless of the mAb-HRP incubation time.

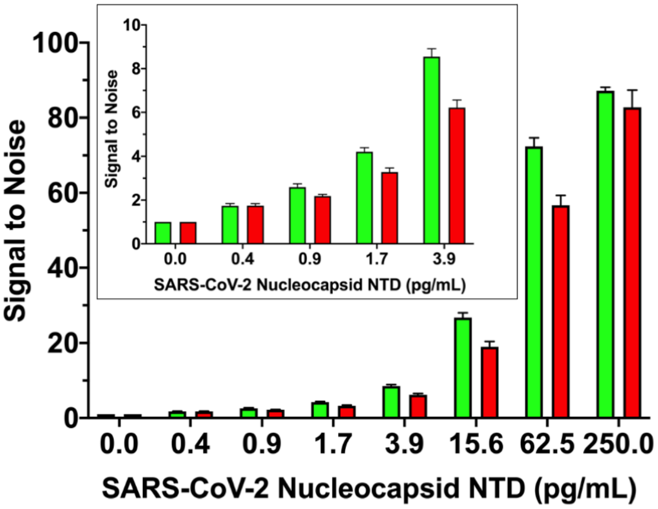


**Supplementary Figure 6. The addition of virion lysing triton X-100 increases the LOD of the modified ELISA, diminishing the signal's boost.**

(A) A modified ELISA where the detection HRP-conjugated mAbs (1-HRP in green, 4-HRP in red) are added directly on top of the samples during the $NP^{NTD}$ capture incubation period, and free (non-plate-bound) mAb2 is “spiked” into the detection HRP-conjugated mAb solutions before their addition on top of the samples. The $NP^{NTD}$ samples were diluted in PBS pH 7.4 plus 0.5% triton X-100.

| Antibody | Association rate constant, $k_a$ ($M^{-1}s^{-1}$) | Dissociation rate constant, $k_d$ ($s^{-1}$) | Dissociation equilibrium constant, $K_D$ (pM) | Experimental Rmax (RU) | Theoretical Rmax (RU) | % Activity of mAb |
|---|---|---|---|---|---|---|
| **mAb 1** | **$1.7 \times 10^{7}$** | **$2.2 \times 10^{-5}$** | **1.3** | **56.4** | **65.0** | **87** |
| **mAb 1-HRP** | **$7.6 \times 10^{6}$** | **$8.5 \times 10^{-5}$** | **11.0** | **13.6** | **45.6** | **30** |
| **mAb 4** | **$1.1 \times 10^{7}$** | **$1.2 \times 10^{-4}$** | **11.0** | **39.4** | **43.4** | **91** |
| **mAb 4-HRP** | **$1.7 \times 10^{7}$** | **$4.7 \times 10^{-4}$** | **28.0** | **4.1** | **40.6** | **10** |
| **mAb 2** | **$9.4 \times 10^{6}$** | **$1.8 \times 10^{-3}$** | **190.0** | **20.7** | **39** | **53** |

**Supplementary Table 1.**
